# A universal pipeline MosaicProt enables large-scale modeling and detection of chimeric protein sequences for studies on programmed ribosomal frameshifting

**DOI:** 10.1101/2025.05.29.656767

**Authors:** Umut Çakır, Noujoud Gabed, Ali Yurtseven, Igor Kryvoruchko

**Affiliations:** Clinical Neuroscience Research Group, Max Planck Institute for Multidisciplinary Sciences, The University of Göttingen, Göttingen 37075, Germany; Cellular and Molecular Biology Department, Oran High School of Biological Sciences (ESSBO), Oran 31000, Algeria; Faculty of Electrical and Electronics Engineering, Istanbul Technical University, Istanbul 34485, Turkey; Department of Biology, United Arab Emirates University, P.O. Box 15551, Al Ain, UAE

**Keywords:** Chimeric peptide, programmed ribosomal frameshifting, alternative open reading frame, proteome, mass spectrometry proteomics, mosaic translation

## Abstract

Peptides and proteins produced by programmed ribosomal frameshifting (PRF) are well-known in viruses. In non-viral systems, only a few examples of such chimeric sequences have been documented until recently. Two new studies, one in humans and one in plants, showed that chimeric peptides are numerous and diverse. In humans, their discovery was possible due to focusing on sequences with naturally repeated codons. This way, many candidate sequences with mass spectrometry (MS) proteomics-based support for translation have been identified. In the plant study, our group discovered MS-validated chimeric peptides using a unique modeling algorithm, which is described and made available here. Our pipeline enables the identification of chimeric peptides in any organism for which transcript sequences and MS proteomic data are available. By design, our approach does not require prior knowledge about sequence similarity to already characterized PRF sites and can detect forward and backward frameshifts by 1 and 2 nucleotides. Thus, our pipeline opens a path for uncovering previously unknown PRF events across various transcript types, potentially broadening our understanding of proteome diversity.

## 1. Introduction

Before 1985, it was assumed that programmed ribosomal frameshifting (PRF) and the resulting chimeric proteins were phenomena unique to viruses. Two studies conducted in *Escherichia coli* were probably the first to suggest that PRF can have a biological role in a non-viral system (Craigen et al., 1985; Craigen and Caskey, 1986). By 2006, it became clear that PRF is found in all kingdoms of life (Dinman, 2006). So far only a few genes have been convincingly shown to use PRF for various purposes in prokaryotes (Craigen et al., 1985; Craigen and Caskey, 1986; Blinkowa and Walker, 1990; Flower and McHenry, 1990; Tsuchihashi and Kornberg, 1990; Chaijarasphong et al, 2016; Meydan et al., 2017) and eukaryotes (Matsufuji et al., 1995; Clark et al., 2007; Ivanov and Atkins, 2007; Ren et al., 2024). Based on bioinformatic predictions, it was proposed that up to 10% of human genes and genes in all eukaryotic genomes can undergo PRF (Belew et al., 2011; Ketteler, 2012; Michel et al., 2012). However, experimental evidence for a high abundance of chimeric peptides in humans was obtained only in 2024. Ren and associates found 405 unique mass spectrometry (MS)-supported chimeric peptides in 32 normal human samples (Ren et al., 2024). These peptides are thought to originate from 454 loci which have naturally occurring repeat codon sequences at the putative PRF sites. Functional characterization of one such locus, which encodes a histone deacetylase HsHDAC1, revealed that the frameshifted product inhibits the non-frameshifted version of HsHDAC1 (Ren et al., 2024). Our parallel study in the model plant *Medicago truncatula* (Çakır et al., 2024, preprint) is conceptually different from the work of Ren et al. (2024). Although it lacks functional validation, we obtained MS support for the translation of 156 chimeric peptides that are not limited to products of mRNA and PRF sites with repeated codons. In addition to mRNA-derived chimeric peptides, we also found that some ncRNA, rRNA, and even tRNA transcripts could potentially produce chimeric peptides. Furthermore, in contrast to the human study, which was focused on PRF events with values - 1 and +1, we extended our scope to include “long” PRF events, encompassing -2 and +2 frameshifts. The discovery of these diverse chimeric peptides in the plant proteome was possible due to our original approach to identifying chimeric peptides via MS, as proposed in our viewpoint article introducing the mosaic translation hypothesis (Çakır et al., 2023). It is based on the identification of conserved and/or translated alternative open reading frames (altORFs) and using them as “building blocks” for chimeric protein models when altORFs overlap with each other and/or with the main annotated ORF (refORF). Translated altORFs and their products, altProts, are thought to be ubiquitous in all organisms, and many of them have been implicated in vital cellular functions (Mouilleron et al., 2016; Orr et al., 2020). A recently developed database called OpenProt provides comprehensive inventory of altORFs in multiple species (Brunet et al., 2019, 2021; Leblanc et al., 2024). Our original pipeline described in this publication was implemented in *M. truncatula*. However, it can be applied to any organism for which transcriptome and MS proteomic data are available. By enabling systematic detection of PRF-derived peptides, this pipeline has the potential to uncover previously unrecognized aspects of proteome diversity and reveal novel biological functions associated with PRF.

## 2. Material and methods

In this manuscript, we use the term “chimeric peptide” for short amino acid sequences deduced from MS proteomics, representing segments produced through PRF. For *in silico* generated models, we use the term “chimeric protein model”. They are hypothetical constructs that match MS-derived chimeric peptides and may represent fragments of longer chimeric or mosaic proteins. When discussing these sequences without specifying length, we refer to them broadly as “chimeric proteins”. Our pipeline consists of three modules that correspond to different steps in the preparation of chimeric protein models: (1) the detection and *in silico* translation of ORFs, (2) the removal of refProts, and (3) the modeling of chimeric proteins based on the validated altProts (Figure 1). The first module identifies ORFs within transcript sequences and performs *in silico* translation to generate potential protein sequences. In the second module, refProts are removed from the list of potential protein sequences deduced by the first module. The third module generates chimeric protein models based on user-defined criteria, e.g., list of validated altProts. These modules are used in conjunction with the software for BLASTP analysis (e.g., DIAMOND, Buchfink et al., 2021) and searching for MS peptide matches (e.g., SearchGUI, Barsnes and Vaudel, 2018; PeptideShaker, Vaudel et al., 2015). These tools help focus on conserved altORFs and match the *in silico* models with MS data, supporting the identification of valid chimeric sequences. The models are generated according to three scenarios, which depend on the location of an altORF relative to its refORF and/or other altORFs (Figure 2). According to the first scenario, an altORF can be completely embedded in its refORF or another long altORF without reaching one of untranslated regions (UTRs). In the second scenario, an altORF can partially overlap with its refORF so that part of the altORF is located in a UTR. In the third scenario, there is no overlap between an altORF and a refORF so that the altORF is fully located within a UTR. Scenario 3 addresses an unconventional situation that may require the forward slippage of ribosomes by more than two nucleotides. It was included in our study (Çakır et al., 2024, preprint) to test whether the translation products of non-overlapping ORFs can be combined in a single continuous polypeptide by frameshifting over longer distances (up to ten nucleotides). This means two ORFs joined in this fashion could belong either to different reading frames or to the same reading frame, in contrast to the other two scenarios. An example of the modeling for the first scenario is shown in Supplementary Figures S1 and S2. Below, we describe each of the three modules.

**Figure 1.**
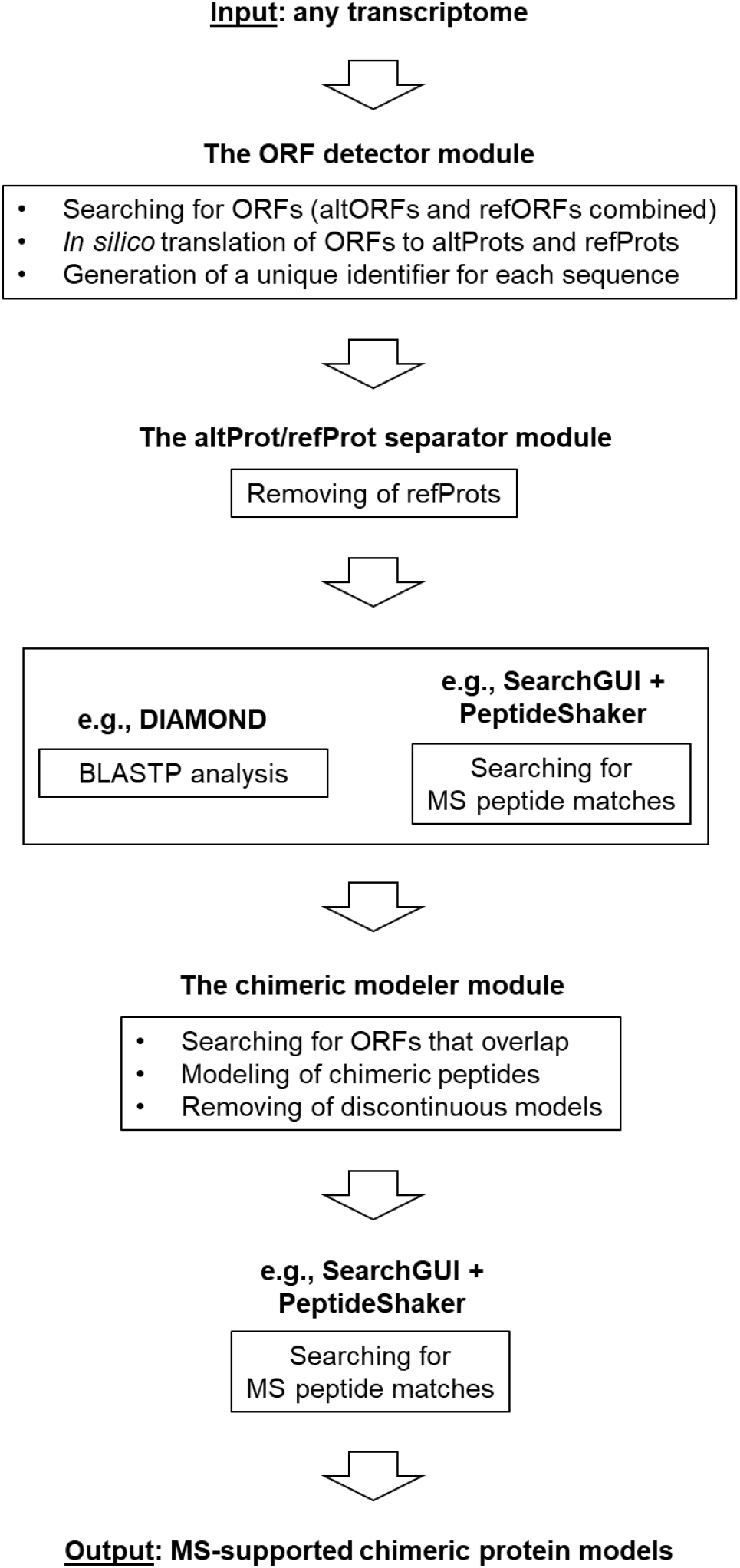
An outline of the detection pipeline for the mass spectrometry (MS)-based discovery of chimeric peptides. Depending on the purpose of analysis, the workflow can start from the chimeric modeler module, which can help find the MS support for chimeric peptides generated from taxonomically restricted ORFs.

**Figure 2.**
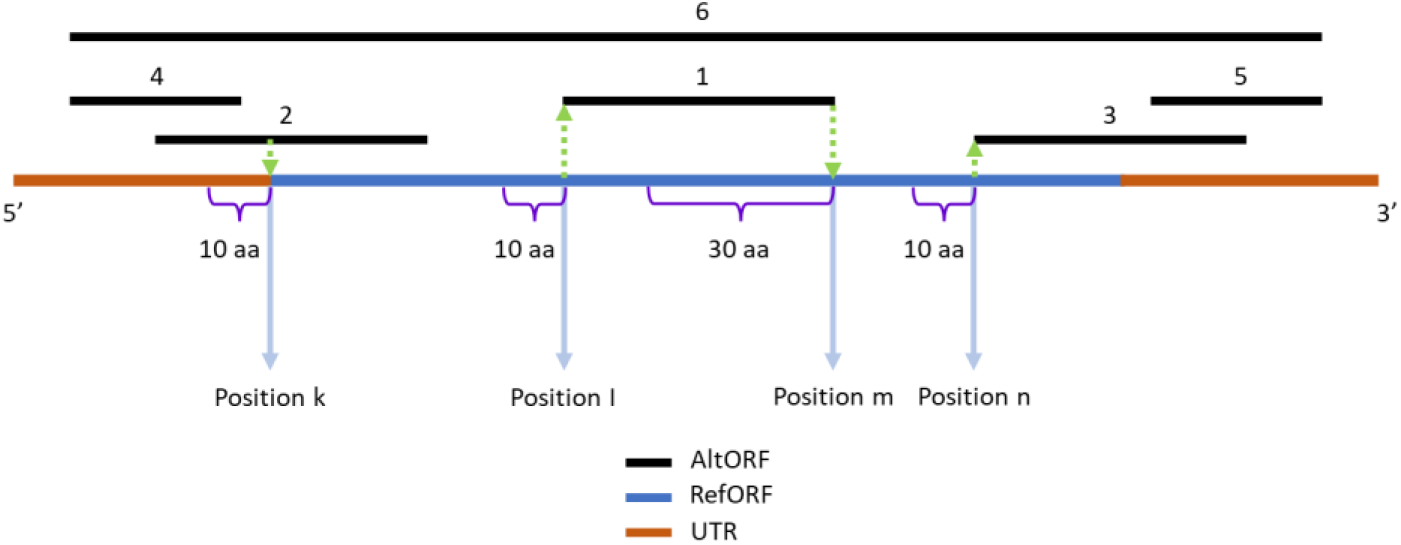
A graphical summary of scenarios handled by the chimeric modeler module. AltORFs and a refORF are shown with black and blue horizontal lines, respectively, and the UTRs are depicted in pale brown. AltORFs can be located in six different places (black numbers above the lines) relative to their refORF, which requires consideration of distinct scenarios. Scenario 1: altORF 1 is embedded in the refORF. Scenario 2: altORF 2 (or altORF 3) overlaps with the refORF. Scenario 3: altORF 4 (or altORF 5) is located entirely in the UTR and does not overlap with the refORF. The shifts from altORF 6 to the refORF (the 5’-side) and from the refORF to altORF 6 (the 3’-side) are considered as special cases of Scenario 2. The identity of positions k, l, m, and n and corresponding distances in amino acids (aa) are explained in the text.

### 2.1. The ORF detector module

The first module identifies and *in silico* translates all ORFs found in a given transcriptome. The module was designed to make no discrimination between refORFs and altORFs. This is an important feature because the status of an ORF as a refORF or an altORF can change over time (Sieber et al., 2018; Wright et al., 2022). In the context of research on PRF using our approach, ORFs are best defined as sufficiently long regions of transcript sequences that are free of in-frame stop codons, regardless of the presence of the canonical start codon AUG. These can be segments of variable length (a user-defined parameter, see Results) found either between two in-frame stop codons (Sieber et al., 2018) or between a stop codon and either end of a transcript. The module provides each generated amino acid sequence with a unique tag (identifier) that contains information about the genomic locus, reading frame, coordinates of an ORF on a source transcript, and the length on the transcript. This detailed tagging process is essential for downstream analyses as it provides clear and accessible information about each ORF, enabling efficient filtering, categorization, and subsequent modeling steps. Users can provide their FASTA-formatted nucleotide sequences (for example, an entire transcriptome) and specify a minimum length threshold for identifying ORFs. By default, the threshold is set to 30 aa (the setting of OpenProt), but users can customize it according to their needs.

### 2.2. The altProt/refProt separator module

The second module is dedicated to filtering out refORF products, or refProts, from the output generated by the first module. A user defines which ORFs are considered as refORFs by providing a FASTA-formatted canonical proteome list. After filtering out refProts, a FASTA file with altProts produced by the module can be used in at least three different ways. Firstly, it can serve as a search database for matching MS peptides found in biological samples, which can provide the most direct evidence for the translation of longer amino acid sequences. We employed SearchGUI v. 4.0.41 (Barsnes and Vaudel, 2018) and its partner tool PeptideShaker v. 2.0.33 (Vaudel et al., 2015) for MS searches. Secondly, the altProt list can be subjected to the global protein sequence similarity search using BLASTP with one of the well-annotated reference protein databases. We used UniProt (UniProt Consortium, 2019) as the database and DIAMOND as a search tool (Buchfink et al., 2021). Such analysis can deliver evidence for the conservation of altORFs at the protein level regardless of the evidence for their translation. Thirdly, the list can be used directly as input for the third module, if the goal is to model chimeric proteins using all possible altORFs above a given length in the entire transcriptome or in its part. The altProt/refProt separator module, therefore, is a critical step in the pipeline, as it refines the dataset to focus on alternative protein-coding sequences. By producing a comprehensive list of altProts, this module enables subsequent analyses that can reveal altORFs’ translational activity, conservation, and potential role in the proteome.

### 2.3. The chimeric modeler module

This module uses four input files to generate chimeric protein models: (1) a transcriptome file, (2) a file that contains altProts, (3) a file that contains refProts, (4) a file that contains a user-defined list of altProt identifiers. Files 2 and 3, which contain altProts and refProts, respectively, are generated by the second module. The fourth file that contains a user-defined list of altProt identifiers may correspond to conserved altProts, MS-supported altProts, or both. Thus, the fourth file may contain a subset of altProt identifiers from the second file. In case users wish to conduct the modeling based on ORFs regardless of their conservation and/or MS support status, the fourth file may either contain altProt identifiers from the entire second file or identifiers of altProts selected based on criteria other than conservation and/or MS support, for example, altProts that originate from one specific chromosome. This module combines three functions: searching, modeling, and filtering. First, it finds overlapping ORFs predefined by input file 4. RefProts involved in the modeling are assumed to be translated. Then, for each overlapping pair of ORFs, it generates a set of chimeric protein models starting at specific positions around the beginning of the overlapping region. The exact starting positions and other parameters are specific to each scenario (Figure 2). The modeling proceeds from left to right in one-nucleotide steps, called iterations, each corresponding to a putative PRF position. PRF values -2, -1, +1, and +2 are considered in our study. The signs correspond to the direction of the frameshift and the number indicates the length of the frameshift in nucleotides. For example, a +2 frameshift advances the ribosome in the forward direction by two nucleotides relative to its normal progression. Our algorithm also addresses special cases where two ORFs do not overlap but are separated by one to ten nucleotides. Translation products of such ORFs can potentially be joined by forward frameshifts with PRF values up to +10. Models generated according to each scenario contain two short amino acid sequences from two reading frames, the left arm and the right arm, with a putative PRF site between them. When the algorithm occasionally produces a model with an empty position that corresponds to an incorporated stop codon, such defective models are removed automatically so that the output file contains only continuous sequences. The final output of the chimeric modeler module is a set of chimeric protein sequences that adhere to the specific PRF scenarios and parameters defined in the study. These models represent theoretical constructs that can be matched with MS data to identify PRF-derived chimeric peptides.

## 3. Theory/calculation

### 3.1. General principles

In this section, we exemplify the calculation of the number of chimeric protein models generated per switch from one ORF to another. It is meant to reflect the computational challenge of including all possible chimeric protein models in the analysis. For the left side of the first modeling scenario (Figure 2, Supplementary Figures S1 and S2), the number (N) of modeled chimeric proteins that correspond to the shift from the refORF (yellow) to the altORF (light red) can be calculated according to Equation 1:

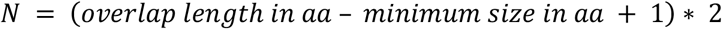

Value “2” in this equation corresponds to making separate models for two possible ways of switching from one frame to another (backward and forward PRF events). Thus, 42 chimeric proteins are modeled for the shift from the refORF to the altORF if the altORF is 90 nt long (30 aa) and if the minimum size of the altORF or the refORF part considered in the modeled chimeric protein is chosen to be 10 aa. In this example, the overlap length corresponds to the whole length of the embedded altORF in amino acids. To illustrate the scale of this process, let us consider a dataset containing 10,000 altORFs, each measuring 90 nt in length and embedded within their respective refORFs. The total number of chimeric models generated for such a dataset can be calculated as follows: 10,000 × 42 for the left side plus 10,000 × 42 for the right side, which makes 840,000 chimeric models. Since many altORF in our study are longer than 90 nt, the actual number of models can be much larger for every increment in the altORF length. This explains why it is practical to consider only a subset of possible models in one MS-based study.

In the first scenario illustrated in Supplementary Figure S1, chimeric proteins can be modeled in two ways that correspond to frameshifts either from the refORF to the altORF or the other way around. In Supplementary Figure S2, chimeric proteins modeled for a frameshift from the refORF to the altORF are shown. Here, if the refORF and the altORF are located in the third frame and the first frame, respectively, the ribosome can switch from the third frame to the first frame in two different ways: via +1 and -2 frameshifting events. There are 42 possible chimeric proteins that can theoretically correspond to the frameshift from the refORF to the altORF if the minimum size of a sequence contributed by either ORF in a chimeric model is chosen to be 10 aa. First, the modeling algorithm takes the region between the nucleotides 390 (10 aa upstream of the altORF start) and 510 (30 aa downstream of the altORF start). Then it proceeds in one-nucleotide steps (iterations) from left to right, moving the frameshift position with each step until the last model contains 30 aa from the refORF and 10 aa from the altORF. For this example, there are 21 iterations and 21 modeled chimeric proteins. The same principle, with some variations, was applied for other scenarios (Figure 2). For the left side of Scenario 1, the iteration process begins 10 aa upstream of the altORF starting point, whereas for the right side, it starts 30 aa upstream of the altORF end. The first model on the right side includes 10 aa from the altORF and 30 aa from the refORF, while the last model contains 30 aa from the altORF and 10 aa from the refORF. The minimum contribution of 10 aa from the first ORF in overlapping pairs is a consistent setting for both sides of Scenario 1. However, in Scenario 2, the last model may include only 1 aa from the second ORF, as this setting accommodates cases such as the copper-related protein in *E. coli*, where a frameshift product included a single amino acid from the alternative frame (Meydan et al., 2017). In Scenario 3, the altORF and refORF do not overlap, but their separation does not exceed 10 nt. Frameshifting events in this scenario involve forward slippage, allowing translation products of non-overlapping ORFs to join. For this scenario, the minimum size of a sequence contributed by each ORF in a chimeric model was set to 20 aa, which means only frameshifting events that join two non-overlapping ORFs were considered. The overall length of each chimeric protein fragment limited to only 40 aa is a parameter common for all the three scenarios. These settings were chosen based on the length range typical for MS-derived peptides (7-35 aa, Swaney et al., 2010). The idea was to make sure an MS peptide spans the frameshift position rather than matches either part of the chimeric model. This, however, does not mean we anticipate the chimeric products to be of that short length only. As we hypothesize, they can be long molecules, but we need to focus on the short fragments corresponding to the frameshift sites in order to validate their chimeric nature.

Below we describe further details of each scenario, corresponding settings, and the calculations. The basis for the chimeric protein modeling algorithm is visualized in Figure 2. Position k corresponds to the beginning of the refORF. Positions l and m correspond to the beginning and the end of altORF number 1, respectively. Position n corresponds to the beginning of altORF number 3. The algorithm focuses on a meaningful subset of all possible situations. For example, starting the iteration process at the end of altORF number 2, the end of refORF, or the middle of altORF number 1 was not considered, which helped alleviate the search database inflation problem (Li et al., 2016; Kumar et al., 2017).

### 3.2. Scenario 1

For the altORFs denoted with number 1 in Figure 2, chimeric proteins are modeled separately at the 5’- and 3’-portion of the altORFs. At the 5’-end of the altORF, the algorithm starts modeling 10 aa upstream of position l and proceeds until position l + 30 aa in 21 iterations. All corresponding chimeric models in this region are generated: the first chimeric protein model is composed of 10 aa from the refORF and 30 aa from the altORF, and the last one is composed of 30 aa from the refORF and 10 aa from the altORF. In contrast, at the 3’-side of the altORF, which is used for the modeling of the switch to the refORF, 30 aa upstream of position m is taken and extended to position m + 10 aa in 21 iterations, so that the last iteration stops at the end of the altORF. All chimeric models in this region are generated: the first model is composed of 10 aa from the altProt and 30 aa from the refProt, and the last model is composed of 30 aa from the altProt and 10 aa from the refProt. The sequence of the first 10 aa is the same for all 21 models in this case. Likewise, the last 10 aa of all 21 models are shared.

### 3.3. Scenario 2

When an altORF that starts in the 5’-UTR overlaps with its refORF (number 2 in Figure 2), chimeric proteins are modeled in the following way: 10 aa upstream of position k are taken and extended via iterations to position k + 30 aa. All chimeric protein models in this region are generated. The first chimeric protein model (the first iteration) is composed of 10 aa from the altProt and 30 aa from the refProt. However, the last chimeric model (iteration 30) is composed of 39 aa from the altProt and only 1 aa from the refProt. All generated chimeric models share the first 10 aa. Their number is calculated as follows: 30 × 2 = 60, where “30” indicates the number of iterations from one to 30, and “2” refers to the separate consideration of backward and forward frameshifting events. When an altORF that ends in the 3’-UTR overlaps with its refORF (number 3 in Figure 2), chimeric proteins are modeled in the following way: 10 aa upstream of position n is taken and extended via iterations to position n + 30 aa. All chimeric models in this region are generated. The first model is composed of 10 aa from the refProt and 30 aa from the altProt. The last model is composed of 39 aa from the refProt and one amino acid from the altProt. Similar to the 5’-UTR side of this scenario, all generated chimeric models share the first 10 aa. Likewise, their number is 30 × 2 = 60.

### 3.4. Scenario 3

When an altORF does not overlap with its refORF, but the gap between the altORF and the refORF is equal to or less than 10 nt (altORFs 4 and 5 in Figure 2), chimeric proteins are modeled in the following way. If the altORF is located at the 5’-UTR (4), the last 20 aa of the altProt and the first 20 aa of the refProt are joined in sequential order. Likewise, if an altORF is located at the 3’-UTR (5), the last 20 aa of the refORF and the first 20 aa of the altProt are joined in sequential order. The number of modeled chimeric proteins is limited to one per each pair of ORFs.

### 3.5. Other scenarios

In addition to the scenarios mentioned above, an altORF may span the whole refORF; that is, the beginning and the end of the altORF can be located at the 5’-UTR and the 3’-UTR, respectively (altORF 6 in Figure 2). In this case, the shifts from altORF 6 to the refORF (the 5’-side) and from the refORF to altORF 6 (the 3’-side) are considered as special cases of Scenario 2, with a few modifications. Namely, 10 aa upstream of position k is taken and extended to position k + 30. While the first chimeric model is composed of 10 aa from the altProt and 30 aa from the refProt, the last chimeric model is composed of 39 aa from the altProt and one amino acid from the refProt. Likewise, the altORF end is handled almost as in the case with altORF 3 with the difference that 30 aa is taken upstream of the refORF stop codon and extended to the total length of 40 aa (30 iterations). The first chimeric model is generated as 10 aa from the refProt plus 30 aa from the altProt. The last chimeric model is composed of 39 aa from the refProt and one amino acid from the altProt. In rare cases, chimeric proteins cannot be modeled as explained above. For instance, if an altORF is shorter than 90 nt and is embedded in its refORF, or if there is no 10 aa long sequence upstream of position k, the length of chimeric protein models can be less than 40 aa. In those rare cases, chimeric proteins can be modeled with the overall length below 40 aa.

Here we explained in detail the overlapping of an altORF with its refORF. However, an altORF can overlap with another altORF. In such cases, one altORF was considered as a refORF, and many possible chimeric models were generated with the same procedure. Furthermore, if more than two ORFs overlap, all paired combinations of ORFs were modeled, and possible chimeric models were generated with the same procedure. For instance, if two altORFs (A and B) and one refORF (C) overlap in the same transcript, the following chimeric models were generated: A & C, B & C, A & B. Here, the ampersand symbol (“&”) denotes the frameshift event that combines products of two overlapping ORFs into a chimeric protein model. By considering all possible pairs, the algorithm ensures that every biologically plausible chimeric protein is captured, accommodating the full complexity of overlapping ORFs in multi-ORF transcripts.

## 4. Results

In our study, we set the minimal length of altORFs to 60 nt while OpenProt uses the threshold of 90 nt (Brunet et al., 2019, 2021; Leblanc et al., 2024). The sequential operation of the ORF detector module and the altProt/refProt separator module identified 875,356 altORFs in the annotated transcriptome v. 5.1.7, which was downloaded from the *M. truncatula* genome portal MtrunA17r5.0-ANR (Pecrix et al., 2018). The transcriptome files that served as inputs for the first module contained transcripts of four types: mRNA, ncRNA, rRNA, and tRNA; 51,317 non-microRNA transcripts in total. Although a precursor of miRNA, pre-miRNA, was shown to act as a template for translation (Couzigou et al., 2016), mature miRNA sequences are too short for the identification of ORFs longer than 59 nucleotides. Thus, they were excluded from the analysis. Excessively large numbers of amino acid sequences included in the search database for the identification of corresponding MS peptides are associated with the phenomenon of database inflation (Li et al., 2016; Kumar et al., 2017). To avoid this problem, in our study, we limited the modeling of chimeric proteins to altORFs that are either MS-supported (805) or conserved (13,078) or MS-supported and conserved at the same time (103); 13,780 unique sequences in total. Different criteria can be used to define what a conserved altORF is. We used a simplified definition of “conserved” by focusing on altORF products that have at least one match to a subject in the UniProt database with a percent identity value of at least 70 and an e-value at most 0.001 in DIAMOND search (Buchfink et al., 2021).

Using 51,317 transcript sequences and 13,780 conserved and/or MS-supported altORFs as inputs, the chimeric modeler module generated 473,871 unique models of chimeric proteins, out of which 472,087 models were built from overlapping ORFs. The remaining 1,784 models were built from non-overlapping ORFs that are not more than 10 nt apart. The lengths of the arms and the overall length of each model were chosen so that the MS peptides cover the PRF sites. This design maximizes the efficiency of detection because the typical length of MS peptides in proteomic studies is 7-35 aa (Swaney et al., 2010). Accordingly, most models are close to 40 aa in length. Among models that were supported with MS proteomics in our study, the length varied between 28 and 41 aa. The length of each arm varied between one and 40 aa. Typically, 42 models were generated per hypothetical shift from one ORF to another, half of which corresponded to backward hypothetical PRF sites (shift by one or two nucleotides from right to left) and the other half contained forward hypothetical PRF sites (shift by one or two nucleotides from left to right). Models generated with such parameters were used as a search database for matching MS peptides present in 16 biological samples generated by three studies (Marx et al., 2016; Shin et al., 2021; Castañeda et al., 2021). Corresponding MS proteomic data were downloaded from PreoteomeXchange (Vizcaíno et al., 2014; Deutsch et al., 2023). The MS searches were conducted independently for models based on conserved and MS-supported altORFs. Likewise, the MS proteomic data from each of the three studies were searched independently. Furthermore, while a regular MS search procedure was applied to non-mRNA-based models and models that come from MS-supported altORFs, a two-step search procedure was used for mRNA-based models that come from conserved altORFs (Jagtap et al., 2013). The same principle was applied during the MS-validation of non-chimeric altProts that were used for the modeling of chimeric events where conserved and mRNA-derived altProts outnumber MS-supported and non-mRNA derived altProts. The purpose of this two-step approach was to alleviate the database inflation effect, which otherwise is expected to increase the rate of false negatives and to compromise the confidence of detection when the search database is large (Li et al., 2016; Kumar et al., 2017). In total, 156 chimeric peptides were supported with MS proteomic data, all of which came from overlapping ORFs. None of 1,784 models generated from non-overlapping ORFs received MS support in our study. Out of 156 MS-supported models, the majority came from conserved altORFs (135). Remarkably, although no rRNA and tRNA-based altORFs were supported with MS (the MS search that preceded the modeling), six chimeric models from rRNA transcripts and one from a tRNA transcript had matching MS peptides. Eleven MS-supported chimeric models corresponded to ncRNA transcripts. The remaining 138 models came from mRNA transcripts. Peptides for the majority of MS-supported models (129) were found in just one sample each, and the remaining 27 models had peptides found simultaneously in two to nine MS samples. Although approximately equal numbers of chimeric models were generated per each PRF value (−2, -1, +1, and +2), sequences that correspond to backward PRF events were significantly overrepresented among MS-supported models. Statistical analysis conducted on these data revealed 37 significant observations and 14 significant associations among various parameters of MS-supported models, their peptides, and corresponding transcripts, which suggests the biological relevance of the dataset (Çakır et al., 2024, preprint).

## 5. Discussion

Although the approach described here was proposed for the *de novo* detection of diverse PRF events in our earlier work (Çakır et al., 2023), to the best of our knowledge, it has not been applied by other groups so far. Presumably, the approach was perceived as too computationally intensive and too prone to false discoveries. In our recent study, we demonstrated that although the approach is tedious, it enabled the discovery of 156 candidates for translated chimeric peptides, some of which correspond to multiple PRF events per transcript (two to three per transcript; 15 non-repeat loci and eight repeat loci). Careful analysis of such multi-PRF sequences revealed two likely candidates for mosaic proteins, which correspond to a putative protein-synthesizing GTPase (a homolog of elongation-factor 1-alpha) and a putative ribulose-bisphosphate carboxylase (RuBisCo). These two completely unrelated loci share the same pattern and parameters of PRF events, which speaks for the biological significance of these shared characteristics (Çakır et al., 2024, preprint). No candidate mosaic sequence is known so far beyond viruses. Thus, our new pipeline generated data that deserve dedicated functional studies, which have the potential for major discoveries, among which the experimental proof of the mosaic translation hypothesis (Çakır et al., 2023) and translation of chimeric peptides from non-mRNA transcripts.

The pipeline presented here is applicable to any organism, providing a tool to identify diverse PRF events without requiring prior knowledge of PRF sequences. This makes our approach uniquely suited to discovering novel PRF sites and events that might evade detection by conventional methods. The pipeline can identify PRF events that join the products of non-overlapping ORFs. With some modifications, the same principle can be applied for the detection of frameshifts “longer” than two nucleotides. In viruses, “long” frameshifts with the slippage of the ribosome by up to six nucleotides have been known for a long time (Weiss et al., 1987; Yan et al. 2015). Although we focused on PRF events with four values (−2, -1, +1, and +2) and have not identified any chimeric peptides that come from non-overlapping ORFs, the sequence diversity of putative PRF sites is enormous. They are almost completely unique to each MS peptide. This emphasizes that our approach can be far more efficient in the search for novel PRF sequences compared to other methods, which can reveal new determinants and mechanisms of PRF in any organism.

As a technical note, modeling chimeric proteins with our pipeline using all ORFs present in a transcriptome would create a database of an astronomic size, which would be difficult to use efficiently for the MS proteomic searches. This is why we limited modeling to conserved and MS-validated altORFs only. However, it may be informative to use altORFs for modeling chimeric proteins regardless of their conservation and/or translation status if the MS search is based on products of only a fraction of the genome, for example, one chromosome or only one chromosome arm. Such approach would not cause the database inflation effect, and it has the potential to identify PRF events that evaded the detection so far. It is known that many translated altProts have either no or very low similarity to annotated proteins (Mouilleron et al., 2016; Orr at al., 2020). This is also evident from the fact that, out of 805 MS-supported non-chimeric altORFs detected in our study, only 103 are conserved. Moreover, the conservation of most altORF is limited to either the *M. truncatula* genome or genomes of other legume plants. Thus, PRF events that involve taxonomically restricted ORFs can be a promising subject for future studies using our pipeline.

## 6. Conclusions

The method and the code described here offer the first approach for the *de novo* identification of PRF events, enabling the systematic and unbiased exploration of frameshifting. It operates independently of prior knowledge about PRF sequences or predefined annotations, facilitating the discovery of both conventional and unconventional PRF events. After our pilot study in *M. truncatula*, we hope that the application of this pipeline in a wide range of organisms will enrich the knowledge about the true diversity of proteomes. Importantly, this knowledge can be the basis for the discovery of mechanisms and phenomena that have remained unnoticed so far.

## Abbreviations

altORF: alternative open reading frame
refORF: reference open reading frame
altProt: alternative protein
refProt: reference protein
PRF: programmed ribosomal frameshifting
mRNA: messenger RNA
ncRNA: non-coding RNA
rRNA: ribosomal RNA
tRNA: transfer RNA
aa: amino acids
nt: nucleotides
MS: mass spectrometry

**Umut Çakır**: Conceptualization, Methodology, Software, Validation, Formal analysis, Resources, Data Curation, Writing - Original Draft, Writing - Review & Editing, Visualization, Funding acquisition. **Noujoud Gabed**: Conceptualization, Validation, Formal analysis, Resources, Writing - Review & Editing. **Ali Yurtseven**, Methodology, Software, Formal analysis. Writing - Review & Editing. **Igor S. Kryvoruchko**: Conceptualization, Methodology, Validation, Formal analysis, Resources, Data Curation, Writing - Original Draft, Writing - Review & Editing, Visualization, Supervision, Project administration, Funding acquisition.

## Data availability

The code for this pipeline, along with instructions on installation and functionality, is accessible from the following GitHub repository and PyPI project:

https://github.com/umutcakir/mosaicprot

https://pypi.org/project/mosaicprot

For convenience, input and output file examples are also provided.

## Acknowledgements

This work was supported by the Scientific and Technological Research Council of Turkey (TÜBİTAK) grants and Boğaziçi University standard research grant (BAP-P) to UÇ and IK (TÜBİTAK 1001 120Z514, TÜBİTAK 1002 120Z247, and BAP-P 18841). Computational analysis was conducted using the server of the Turkish National e-Science e-Infrastructure (TRUBA) center. The completion of this study was possible due to the support of IK by United Arab Emirates University and the support of UÇ by the IMPRS-Genome Science PhD program.

## Supporting information

This preprint contains supplementary materials: Supplementary Figures (2). They are available as separate files.

**<u>Supplementary Figure S1</u>**. Generation of model sequences for frameshifted proteins translated around a candidate programmed ribosomal frameshifting (PRF) site is a computational challenge.

**<u>Supplementary Figure S2</u>**. A modeling example for the left-hand side of Supplementary Figure S1.

