## Supplementary Figure S1 for "A universal pipeline MosaicProt enables large-scale modeling and detection of chimeric protein sequences for studies on programmed ribosomal frameshifting"

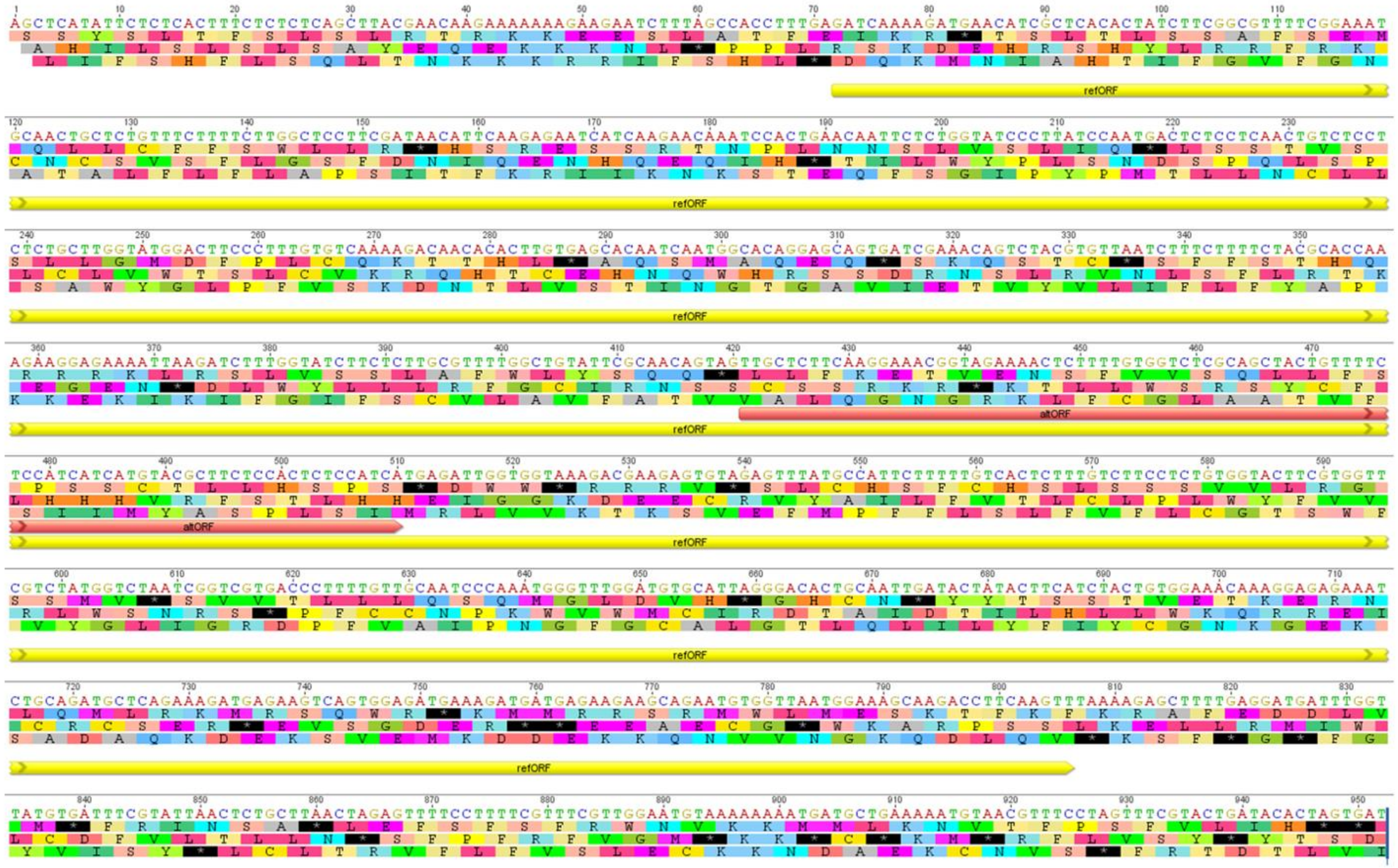


**Supplementary Figure S1**. Generation of model sequences for frameshifted proteins translated around a candidate programmed ribosomal frameshifting (PRF) site is a computational challenge. This image illustrates a hypothetical transcript and its three-frame translation. There is one reference open reading frame (refORF, yellow) in frame 3 and one translated alternative open reading frame (altORF, light red) in frame 1 that overlaps the refORF. The exact location of the hypothetical PRF event is unknown.
