## Supplementary Figure S2 for "A universal pipeline MosaicProt enables large-scale modeling and detection of chimeric protein sequences for studies on programmed ribosomal frameshifting"

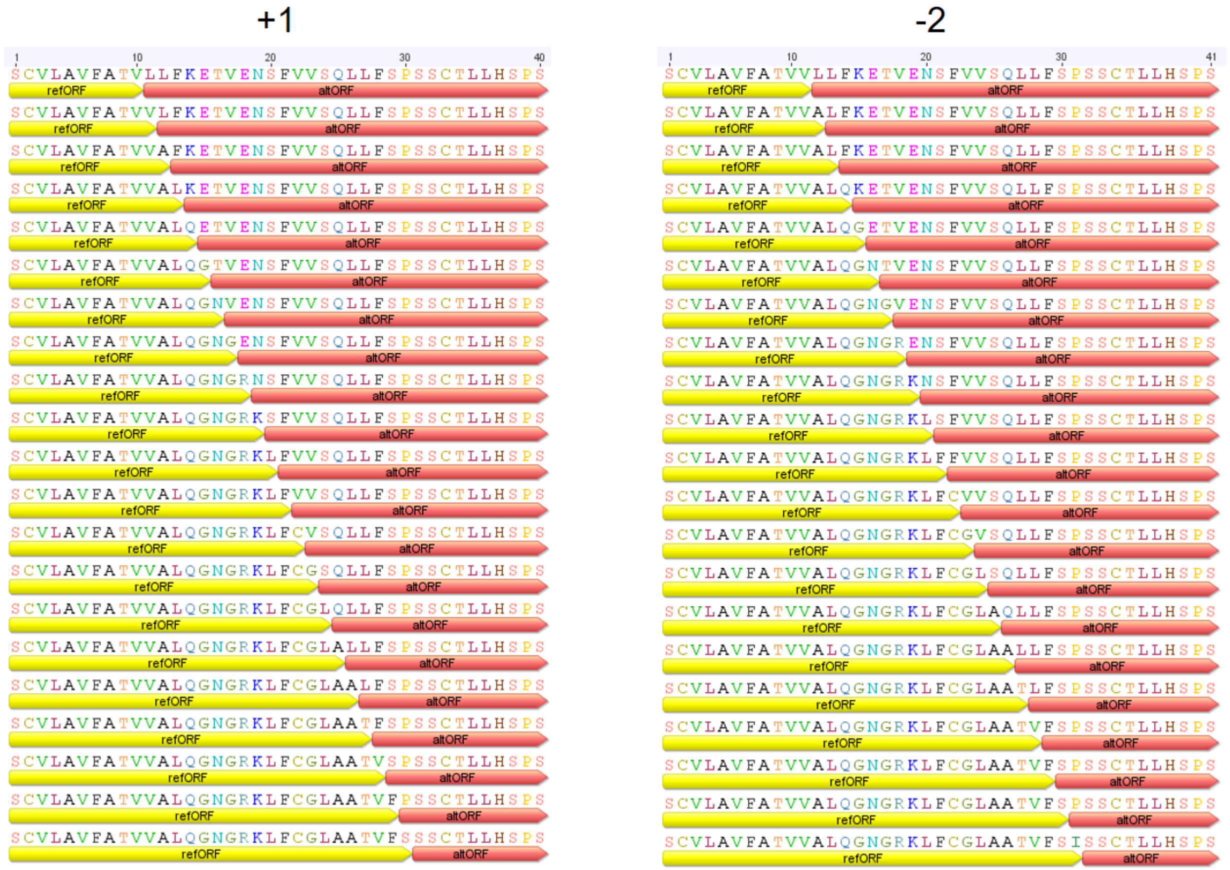


**Supplementary Figure S2**. A modeling example for the left-hand side of Supplementary Figure S1. There are 42 chimeric proteins that can theoretically correspond to the shift from the refORF to the altORF of a certain length (30 aa in this example). “42” comes from the following calculation: (30 - 10 + 1) x 2, or more generally (overlap length - 10 + 1) x 2. “10” is the minimal size of an altORF product in a chimeric protein (an arbitrarily set threshold convenient for the reliable matching to MS peptides on either side of the shift). The left column depicts chimeric proteins (partial view, only the hypothetical PRF site shown) resulting from skipping one nucleotide (PRF value +1). The right column lists proteins resulting from a backward shift by two nucleotides (PRF value -2).
